# biomeStat: Using Agentic AI for Scalable Genomic Epidemiology Demonstrated Through End-to-End Analysis of 1,000 Asian Dengue Virus Genomes

**DOI:** 10.64898/2026.06.10.731380

**Authors:** Dinuka Ariyaratne, Nayana Somaratna, Gathsaurie Neelika Malavige

## Abstract

Genomic epidemiology workflows typically require expert curation of multiple specialized tools, extensive manual parameter tuning, and access to heterogeneous compute infrastructure. While standard generative AI models often hallucinate in complex biological domains, we introduce biomeStat: an autonomous AI agent that functions as a strict deterministic orchestrator. By automatically writing code to execute established bioinformatics tools in sandboxed environments, biomeStat dynamically provisions compute resources (CPU and GPU) and guarantees reproducibility, making it immediately useful for scientists without requiring command-line expertise.

To demonstrate the platform, we performed a fully autonomous genomic epidemiology and structural analysis of 1,000 Dengue virus (DENV) genomes sampled from 16 Asian countries between 2000 and 2025. The agent seamlessly orchestrated phylogenetic reconstruction (IQ-TREE, TreeTime), Bayesian phylodynamics (BEAST2 via NVIDIA H200 GPU), selection pressure analysis (HyPhy), and structural mapping (PyMOL). The analysis was completed in under 24 hours of wall-clock time, revealing endemic stability (R_e ∼1.0) and identifying 1,869 candidate immune escape sites structurally colocalized with B-cell and T-cell epitopes. Furthermore, the agent validated 176 highly conserved drug target residues across the viral replication complex, confirming that resistance-associated positions for emerging antivirals JNJ-1802 and NITD-688 remain absolutely conserved across all four serotypes. By bridging the gap between natural language intent and deterministic computational execution, biomeStat reduces weeks of expert effort into a single-session analysis with full methodological transparency.

## Introduction

Genomic epidemiology is at the core of how we track viral evolution, infer transmission networks and monitor immune escape[1]. However, the actual task of applying phylogenetic and phylodynamic models to emerging viruses such as dengue, SARS-CoV2, influenza etc., faces severe computational and bioinformatics bottlenecks [2]. This chokepoint is often worse in resource constrained research centres especially in the Global South [3–5]. Traditional Bayesian phylogenetic tools, which provide the most accurate epidemiological insights, scale exponentially with dataset size (O(N2)), rendering them computationally intractable for massive datasets like the millions of genomics sequenced during the COVID19 pandemic [2, 6]. This scalability crisis forces researchers to either wait weeks for convergence or rely on heuristic approaches that sacrifice statistical certainty.

Beyond sheer data volume, complex evolutionary dynamics and uneven sampling compound these computational hurdles. For example, early in the SARS-CoV-2 pandemic, the virus’s low genetic diversity led to “flat trees” with unresolved polytomies, making transmission inference mathematically difficult [6]. Furthermore, events like recombination in dengue and SARS-CoV-2 variants, or reassortment in influenza, violate the assumptions of strict bifurcating trees and require computationally heavy network or structured coalescent models [7]. Additionally, unstandardized amplicon sequencing pipelines had introduced systematic errors into millions of SARS-CoV-2 consensus sequences in public databases, generating spurious phylogenetic branches that corrupted downstream evolutionary analyses and required reprocessing of 4.47 million genomes through a purpose-built amplicon-aware pipeline to correct [8]. In the field, real-time sequencing of outbreaks like Ebola and Zika generates data that demands resource-intensive error-correction algorithms simply to distinguish sequencing noise from true mutations [9, 10]. Consequently, executing a complete genomic epidemiology workflow requires integrating numerous specialized tools, extensive parameterization, and managing disparate computational resources (from CPU tasks to GPU-intensive MCMC chains), which typically demands weeks of expert bioinformatician effort[2]. The scale of this challenge was recently underscored when systematic amplicon-driven errors were discovered to have corrupted millions of SARS-CoV-2 phylogenies globally, requiring reprocessing of 4.47 million genomes with purpose-built assembly tools[8].

Here we introduce biomeStat (www.biomestat.com), an AI agent designed to autonomously execute bioinformatics workflows by selecting appropriate tools, configuring parameters, interpreting intermediate results, and dynamically scaling compute resources. The agent made real-time decisions about tool selection, parameter optimization, error handling, and resource allocation without human intervention during execution.

To validate this approach, we tasked biomeStat with a complete genomic epidemiology analysis of 1,000 DENV genomes from Asia spanning 25 years (2000–2025). The workflow produced phylodynamic estimates of effective reproductive number (*R_e_*), serotype-specific demographic histories, a comprehensive map of immune escape candidates, and identification of conserved drug targets, all within a single analysis session.

We selected the dengue virus (DENV) to demonstrate the capabilities of the biomeStat platform because it exemplifies the transition of a historically neglected tropical disease into an urgent global health threat.

Together, the mapped escape sites, conserved antiviral targets, and phylodynamic estimates provide actionable inputs for vaccine design, antiviral development, and public health planning [58].

## Materials and Methods

### Dataset Assembly

We retrieved 1,000 complete DENV genome sequences from NCBI GenBank using the Entrez API. The authors set the sequence curation criteria as: complete genomes, all serotypes, from Asia, collected from human samples, between 2000-2025.

Sequences were stratified to ensure geographic and temporal representativeness: a maximum of 200 sequences per country across 16 Asian nations (Vietnam, *n* = 200; Singapore, *n* = 172; China, *n* = 159; Cambodia, *n* = 91; Philippines, *n* = 86; Thailand, *n* = 68; Indonesia, *n* = 60; India, *n* = 58; Sri Lanka, *n* = 36; Bangladesh, *n* = 25; Pakistan, *n* = 20; Malaysia, *n* = 10; Myanmar, *n* = 5; Japan, *n* = 5; Taiwan, *n* = 4; Laos, *n* = 1)[11]. Temporal coverage spanned 2000–2024, with peak representation in 2006–2008 and 2015–2019. Serotype composition was: DENV-1 (*n* = 371, 37.1%), DENV-2 (*n* = 304, 30.4%), DENV-3 (*n* = 235, 23.5%), and DENV-4 (*n* = 90, 9.0%).

### Lineage Assignment

Serotype-specific lineage classification was performed using Nextclade v3 [11] with reference datasets for each serotype. Lineages were assigned following the nomenclature proposed by Hill et al. [11], which provides a four-level hierarchy (serotype, genotype, major lineage, minor lineage) analogous to the Pango system used for SARS-CoV-2.

### Multiple Sequence Alignment

Whole-genome alignment was performed using MAFFT v7 [12] with default parameters. To study specific viral proteins (Envelope, NS1, NS3, NS4A, NS4B and NS5), we first identified their exact locations within each serotype’s genome. We then extracted these specific genetic sequences, translated them into amino acids, and aligned them using the highly accurate MAFFT L-INS-i algorithm[12].

### Maximum Likelihood Phylogeny

Phylogenetic inference was performed using IQ-TREE 2 [13] under the GTR+F+G4 substitution model [14, 15] with 1,000 ultrafast bootstrap replicates [16]. The analysis yielded a tree with log-likelihood of −359,017, total tree length of 7.55 substitutions/site, and 56.2% of branch length internal to the tree, indicating well-resolved deep structure.

### Molecular Clock and Temporal Analysis

Root-to-tip regression and time-scaled phylogeny construction were performed with TreeTime[17], yielding an estimated substitution rate of 9.16 × 10^−5^ substitutions/site/year across the full dataset.

### Bayesian Phylodynamics

#### Birth-Death Skyline (BDSKY)

A Birth-Death Skyline model [18] was implemented in BEAST2 v2.7.8 [19] on the full 1,000-taxon dataset. The MCMC chain was run for 10^7^states with five-time intervals (2000–2005, 2005–2010, 2010–2015, 2015–2020, 2020–2025). GPU acceleration was provided by the BEAGLE library [20] running on an NVIDIA H200 GPU (141 GB VRAM). Convergence was assessed by effective sample size (ESS) criteria.

#### Per-Serotype Bayesian Skyline Plot (BSP)

Serotype-specific demographic reconstructions were performed using the Bayesian Skyline Plot coalescent model within BEAST2 for each of the four serotypes independently (DENV-1: 371 taxa; DENV-2: 304 taxa; DENV-3: 235 taxa; DENV-4: 90 taxa). Each MCMC chain was run for 10^7^states under an HKY+G4 model with a strict molecular clock. Population size was estimated in five intervals per serotype. A 10% burn-in was applied to all posterior samples.

#### Selection Pressure Analysis

Site-specific selection was quantified using the HyPhy package. Pervasive selection was estimated by FUBAR (Fast Unconstrained Bayesian Approximation for inferring selection) [21], and episodic diversifying selection was detected by MEME (Mixed Effects Model of Evolution) [22]. Additionally, within-serotype Tajima’s D[23] was computed for E, NS1, NS3,NS4, and NS5 to characterize population-level selection.

#### Immune Escape Mapping

To predict where the virus is actively mutating to evade the immune system, we mapped candidate immune escape sites by integrating structural, immunological, and evolutionary data across the viral proteins.

To structurally visualize population-level sequence variability, position-specific Shannon entropy values calculated from the 1,000-genome multiple sequence alignment were mapped onto the 3D crystal structure of the DENV-2 Envelope (E) glycoprotein complexed with the broadly neutralizing human antibody EDE1 C10 (PDB ID: 4UT9). Similarly, to map T-cell immune evasion, predicted MHC Class I and II epitopes and episodic escape mutations were mapped onto the structure of the NS5 RNA-dependent RNA polymerase and Methyltransferase domains (PDB ID: 6IZZ). Using PyMOL (Schrödinger, LLC), entropy values were substituted into the temperature factor (B-factor) column of the PDB files to render surface variability, allowing direct spatial correlation between mutational hotspots, immune epitopes, and functionally constrained viral domains.

Rather than randomly analyzing mutations across the entire genome, we restricted our search to regions most likely to be targeted by the host immune response. To avoid the severe computational bottleneck of querying external machine-learning web servers for 1,000 full-length genomes, we adapted the underlying biological logic of these platforms [24, 25] for rapid, offline analysis:

We identified B cell epitopes by computationally analyzing the inherent physicochemical properties required for antibody binding. Specifically, we calculated local hydrophilicity using the Hopp-Woods scale[26] and predicted surface accessibility using the Emini fractional probability model [27]. By establishing strict statistical thresholds for these metrics, we effectively flagged surface-exposed, water-accessible regions, accurately mirroring the logic of platforms like BepiPred 3.0[24, 28].

To find potential T cell epitopes, MHC Class I and Class II presentation sites were identified by applying the binding affinity rules derived from the NetMHCpan 4.1 and NetMHCIIpan 4.0 neural networks[25]. We computationally scanned our viral alignments to locate conserved anchor residues and sequence profiles that satisfy the strong-binding definitions (e.g., predicted binding rank ≤2%) established by these models. This allowed us to map the boundaries of peptides most likely to be naturally presented by diverse human leukocyte antigen alleles [29, 30].

To measure how much these specific immune-targeted regions mutate across our Asian Dengue isolates, we calculated the Shannon entropy [31]at every single amino acid position. High entropy flags a hotspot where the virus actively tolerates or encourages amino acid variability, whereas low entropy indicates a structurally conserved residue.

When identifying true escape mutants because high variability alone does not prove immune escape (it could simply be random genetic drift), we required proof of evolutionary adaptation. Because absolute cutoff values fail to capture the extreme structural variability between different Dengue proteins, we implemented a dynamic, protein-specific heuristic. We operationally defined a 70th percentile composite propensity score and a 6-residue minimum length constraint because absolute thresholds fail across highly heterogeneous viral proteins.

This dynamic, protein-specific threshold successfully flagged the most structurally viable antibody targets. Crucially, any potential false positives generated by this structural screening were subsequently filtered out by our rigorous evolutionary criteria (FUBAR/MEME) [21], ensuring that only sites under active positive selection were ultimately classified as immune escape candidates.

We identified true immune escape sites by looking for a specific signature: when a mutation inside an epitope was both highly variable across strains and statistically proven to be under positive selection, we concluded the virus was actively retaining this mutation to evade host immune defenses.

### Drug Target Identification

Conserved functional domains in NS3 (protease catalytic triad, helicase Walker motifs, DECH box) and NS5 (STAT2-binding interface, RdRp GDD motif) [32–34]were evaluated for conservation across all 1,000 genomes. Conservation was quantified as the fraction of sequences with the consensus residue at each position. Positions with >99.5% conservation, catalytic function, and strong purifying selection were classified as high-confidence drug targets[32, 33, 35]. To assess conservation at the binding sites of emerging NS4B-targeting antivirals, we also the NS4A (125 amino acids) and NS4B (250 amino acids) proteins from all genomes Drug resistance-associated positions (NS4B P104, T108 and A119, DENV-2 numbering)[36, 37] were evaluated across all four serotypes independently.

### Codon Usage Analysis

Because dengue virus must replicate efficiently in two drastically different environments namely, the *Aedes aegypti* mosquito vector and the human host, it faces conflicting evolutionary pressures on its genome[38].

To evaluate how the virus balances these conflicting demands, we calculated the Codon Adaptation Index [39] relative to both human and *Aedes aegypti* codon usage tables along with the Effective Number of Codons (ENC) [40] to quantify the degree of host-specific genetic adaptation. CpG dinucleotide ratios were also calculated per genome[41], as the suppression of CpG motifs is a known mechanism by which RNA viruses evade host innate immune surveillance [42].

## Results

### Geographic and Temporal Sampling

The 1,000-genome dataset provided broad coverage of Asian DENV diversity, spanning 16 countries across South, Southeast, and East Asia over a 25-year window (Table 1). DENV-1 was the most represented serotype (37.1%), consistent with its dominance in several Southeast Asian countries during the study period. DENV-4, was the least represented (9.0%).

**Table 1.** This table shows the actual execution time achieved by the biomeStat AI Agent (utilizing dynamic CPU-to-GPU escalation and autonomous orchestration).

| Workflow Phase / Tool | Task Description | biomeStat AI Agent (Autonomous + Scaled) |
| --- | --- | --- |
| Data Curation & QC<br>(Entrez, Nextclade) | Downloading 1,000 genomes, filtering by metadata, lineage assignment | 30 minutes<br>(Automated API calls) |
| Multiple Sequence Alignment<br>(MAFFT L-INS-i) | Full-genome alignment, splitting into polyprotein coordinates (E, NS1, NS3, NS5) | 5 minutes<br>(CPU Tier: 24 vCPU) |
| Phylogenetic Inference<br>(IQ-TREE 2) | Model selection (GTR+G4), ML tree construction, 1,000 UFBoot replicates | 20 minutes<br>(CPU Tier: 24 vCPU) |
| Molecular Clock & Temporal Signal<br>(TreeTime) | Root-to-tip regression, global substitution rate estimation | 10 minutes |
| Selection Pressure & Codon Usage<br>(HyPhy, CAI/ENC) | FUBAR, MEME, Tajima's D, codon adaptation relative to human/mosquito hosts | 15 minutes |
| Immune Escape Mapping<br>(Surrogate Algorithms) | Calculating hydrophilicity, surface accessibility, entropy across 2,365 sites | 10 minutes |
| BEAST2 XML Setup & Debugging | Configuring BDSKY & BSP parameters, resolving namespace/operator bugs | < 1 minute<br>(Autonomous error correction) |
| Bayesian Phylodynamics<br>(BEAST2 MCMC) | Full BDSKY (10M states) + 4 per-serotype BSP chains (10M states each) | 18.4 hours<br>(GPU Tier: NVIDIA H200, autonomous BEAGLE config) |
| Data Synthesis & Visualization | Parsing MCMC logs (Re, Ne), annotating trees, generating 8 publication-quality figures | 5 minutes<br>(Automated Python scripts) |
| Structural Mapping (PyMOL) | 3D surface mapping of Shannon entropy, T-cell epitopes, and functional domains onto PDB 6IZZ (NS5) | 5 minutes (Automated Python/PyMOL scripts) |
| Total Wall-Clock Time | End-to-End Genomic Epidemiology Workflow (1,000 genomes) | ~19.5 Hours |

### Phylogenetic Structure

The ML phylogeny resolved four monophyletic serotype clades with high bootstrap support (>95% for all serotype-level splits). The tree exhibited strong temporal signal, with root-to-tip regression yielding an *R*^2^ of 0.85 against sampling dates. Internal branches comprised 56.2% of total tree length, indicating substantial shared evolutionary history and well-resolved deep branching.

### Bayesian Phylodynamics

#### Serotype-Specific Demographics

Per-serotype BSP analyses revealed distinct effective population size (*N_e_*) trajectories (Figure 3A–D). DENV-2 showed the highest *N_e_* peaks (median 306 in interval 3), consistent with its known capacity for rapid epidemic expansion. DENV-1 showed a peak in interval 2 (*N_e_* = 189) followed by contraction, potentially reflecting lineage turnover events. DENV-4, with the smallest sample (*n* = 90), showed the most uncertainty but a late-period *N_e_* increase.

**Figure 1:**
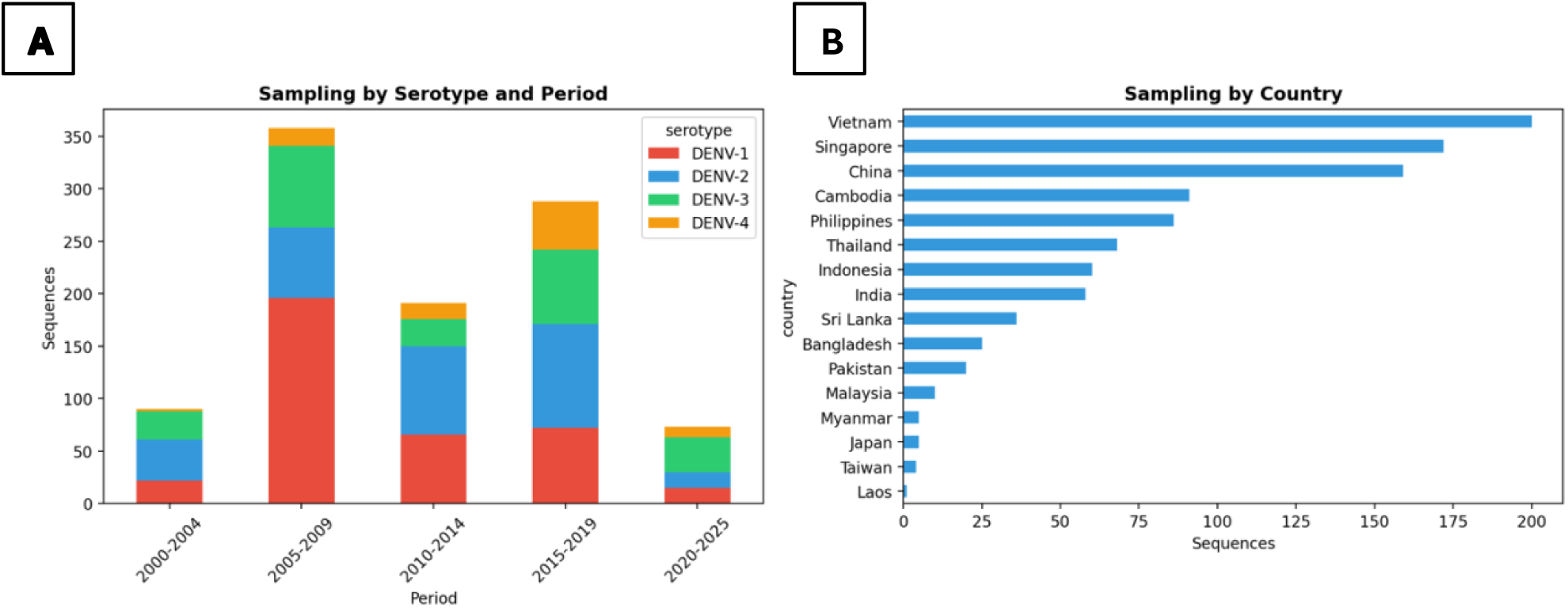
Geographic and Temporal Distribution of the 1,000 Dengue Genomes Description: (**A**): sampling by dengue serotypes and 5-year period spanning 25 years. (**B**): The number of samples per country

**Figure 2.**
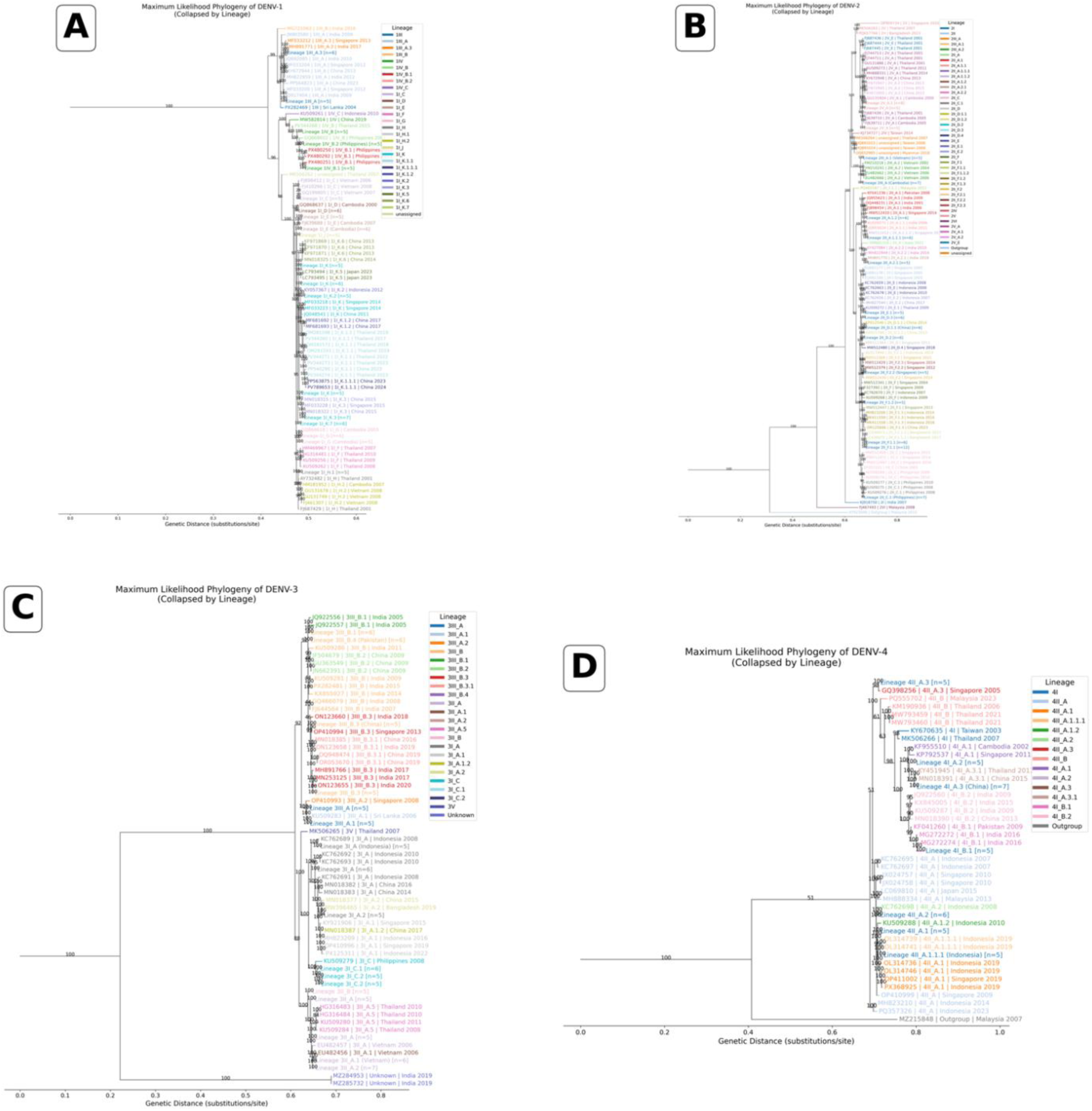
Maximum likelihood phylogenies of 1,000 Asian Dengue virus genomes (2000–2025). Trees were inferred for the four co-circulating serotypes: (**A**) DENV-1, (**B**) DENV-2, (**C**) DENV-3, and (**D**) DENV-4. To enhance readability, highly homogeneous monophyletic clades comprising five or more sequences from the same genetic lineage and geographic region were collapsed into representative nodes. Collapsed nodes are annotated by their specific sub-lineage (following the Hill *et al.* 2024 nomenclature)[11], the dominant country of sampling, and the number of collapsed sequences [n]. Uncollapsed tips are labeled with their accession number, assigned lineage, country, and year of collection. Branch colors and terminal labels correspond to the assigned genetic lineages within each serotype, with the respective color keys provided in the top-left of each panel. The scale bar indicates genetic distance in substitutions per site.

**Figure 3:**
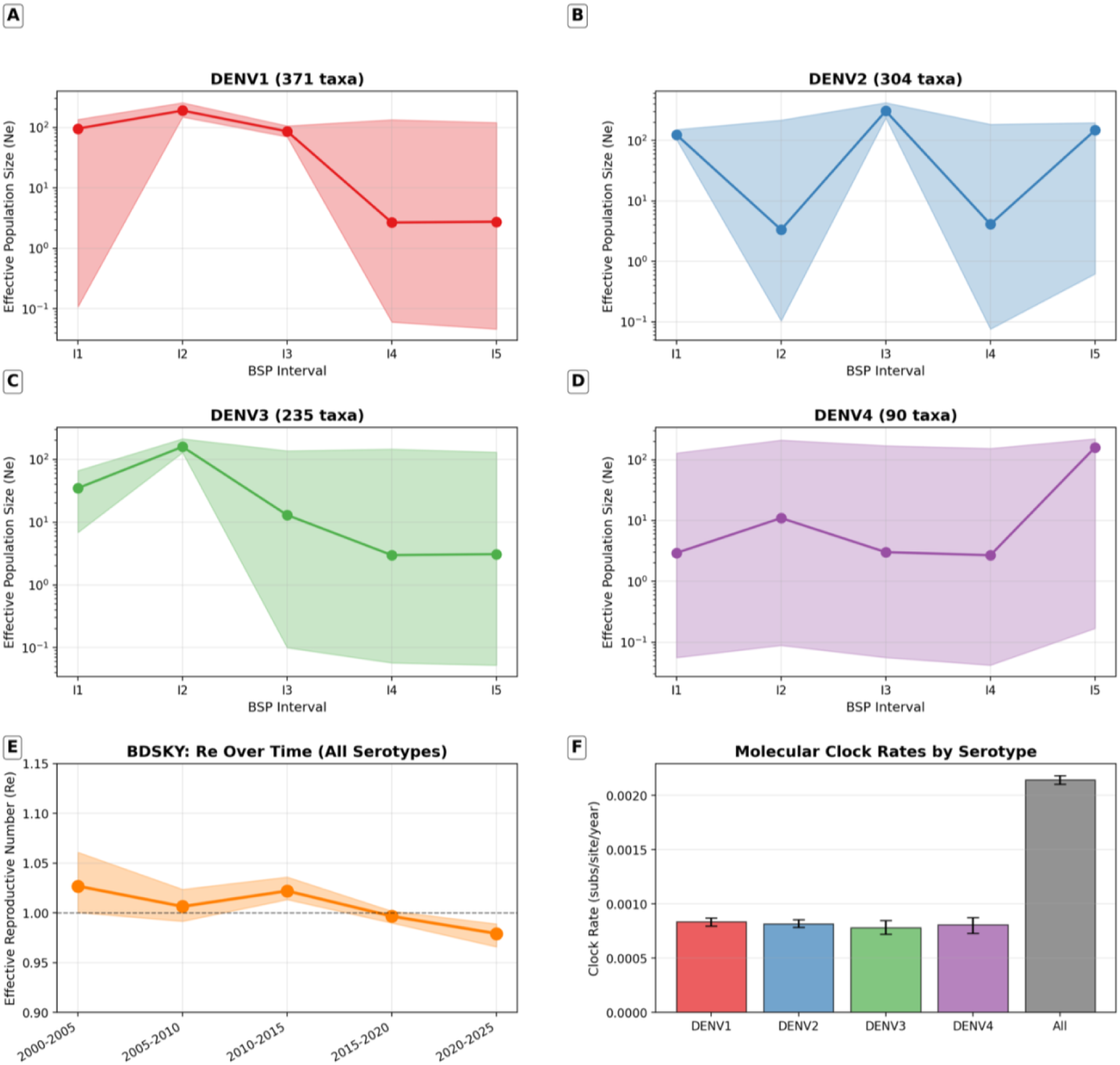
25-Year Epidemiological History and Transmission Dynamics (BEAST2) Description: A summary of the AI-orchestrated BEAST2 phylodynamic modeling. **A–D:** Show the estimated historical population size (number of actively transmitting viruses) for each of the four serotypes. Peaks represent major historical epidemic waves. **E:** Tracks the Effective Reproductive Number (R*_e_*) over five-year intervals from 2000 to 2025. The line hovers closely around the 1.0 threshold. **F:** Compares the molecular clock rate across the serotypes, showing a consistent speed of roughly 8 mutations per 10,000 sites per year.

#### Effective Reproductive Number

The BDSKY analysis estimated *R_e_* across five intervals (Figure 3E). Values oscillated narrowly around 1.0: *R_e_* = 1.027 (95% HPD: 1.001–1.061) for 2000–2005, *R_e_* = 1.007 (0.992–1.024) for 2005–2010, *R_e_* = 1.022 (1.014–1.036) for 2010–2015, *R_e_* = 0.997 (0.990–1.002) for 2015–2020, and *R_e_* = 0.979 (0.966–0.989) for 2020–2025.

#### Molecular Clock Rates

Serotype-specific clock rates were remarkably consistent: DENV-1 (8.38 × 10^−4^), DENV-2 (8.19 × 10^−4^), DENV-3 (7.82 × 10^−4^), and DENV-4 (8.09 × 10^−4^) subs/site/year(Figure 3F). The per-serotype BSP clock rates were approximately 10-fold higher than the full-dataset TreeTime rate (9.16 × 10^−5^). which is expected: TreeTime estimated rates across inter-serotype divergence (deeper time, more saturation), while per-serotype BSP rates reflect intra-serotype evolution over shorter timescales, which is less affected by multiple-hit saturation.

### Selection Pressure

Tajima’s D was strongly negative across all protein-serotype combinations (Figure 4), indicating population expansion and/or purifying selection. Values ranged from −0.86 (NS4A, DENV-2) to −2.53 (NS5, DENV-2). The NS5 protein consistently showed the most negative Tajima’s D values (DENV-1: –2.40; DENV-2: –2.53; DENV-3: –2.45). NS4A in DENV-2 recorded the least negative score (–0.86), followed by NS1 in DENV-4 (–1.21). Overall, E and NS1 proteins (for DEN1,2 &4 serotypes) showed fewer negative values compared to NS5.

**Figure 4:**
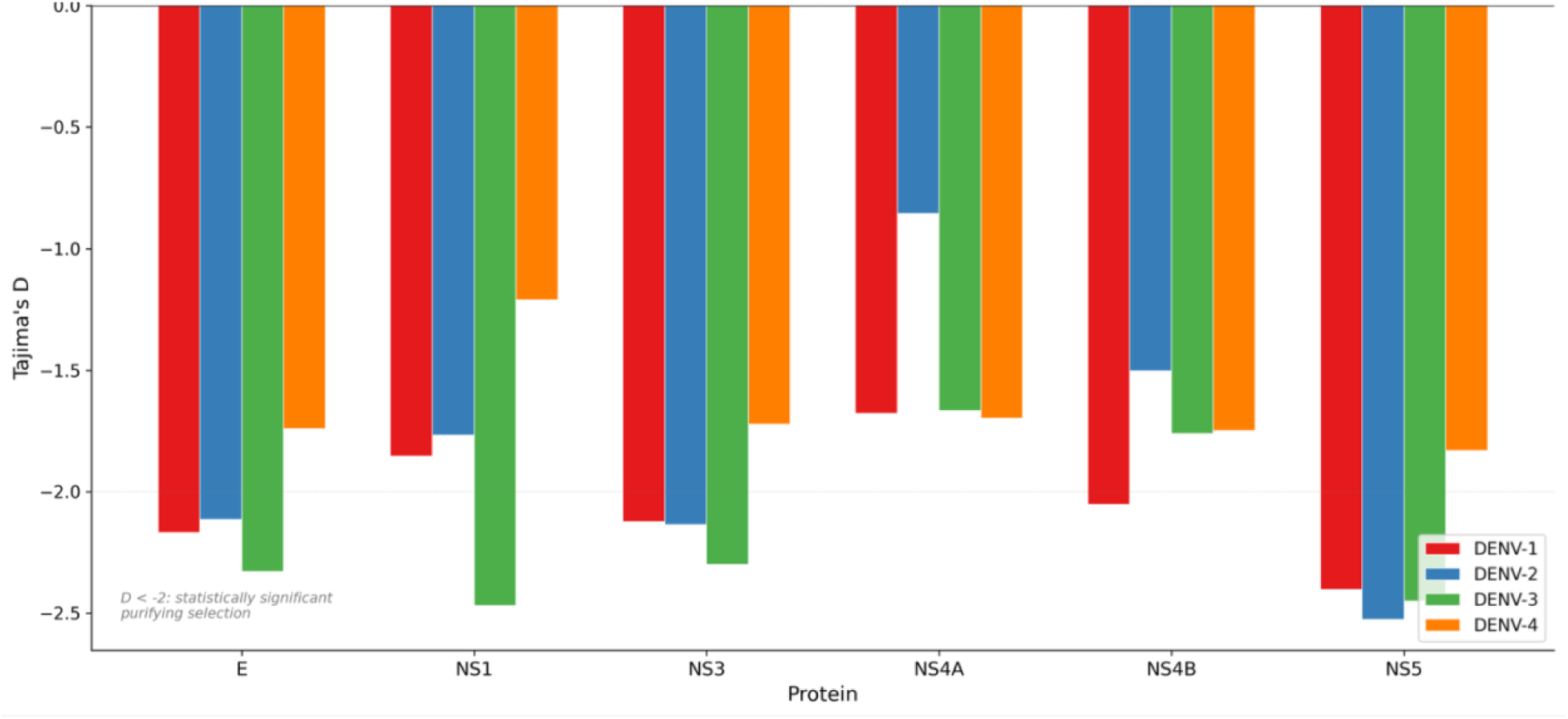
Natural Selection Pressures (Tajima’s D) by Serotype Description: Tajima’s D, calculated for each protein (E, NS1, NS3, NS4A, NS4B & NS5) and each of the four Dengue serotypes. All values fall into the negative range (below zero). D<-2 statistically significant purifying selection.

### Immune Escape Landscape

Of the 1,000 curated DENV1-4 genomes, between 991 and 993 yielded intact open reading frames for each of the six proteins analyzed (E: 993; NS1: 993; NS3: 992; NS4A: 992; NS4B: 992; NS5: 991), with the minor attrition reflecting isolated truncations or frameshifts in individual polyprotein regions.

Of 2,955 protein sites analyzed across E, NS1, NS3, NS4A, NS4B, and NS5, 1,869 (63.2%) were classified as candidate immune escape sites based on the joint criteria of epitope location, elevated entropy, and evidence of diversifying selection. To validate these candidates, we cross-referenced their positions against the 9,909 experimentally validated linear B-cell and T-cell epitopes retrieved from the Immune Epitope Database (IEDB). Strikingly, out of the 1,869 computationally identified escape candidates, 1,355 positions (72.5%) fell directly within a perfectly sequence-matched, experimentally validated IEDB epitope (Figure 5).

**Figure 5:**
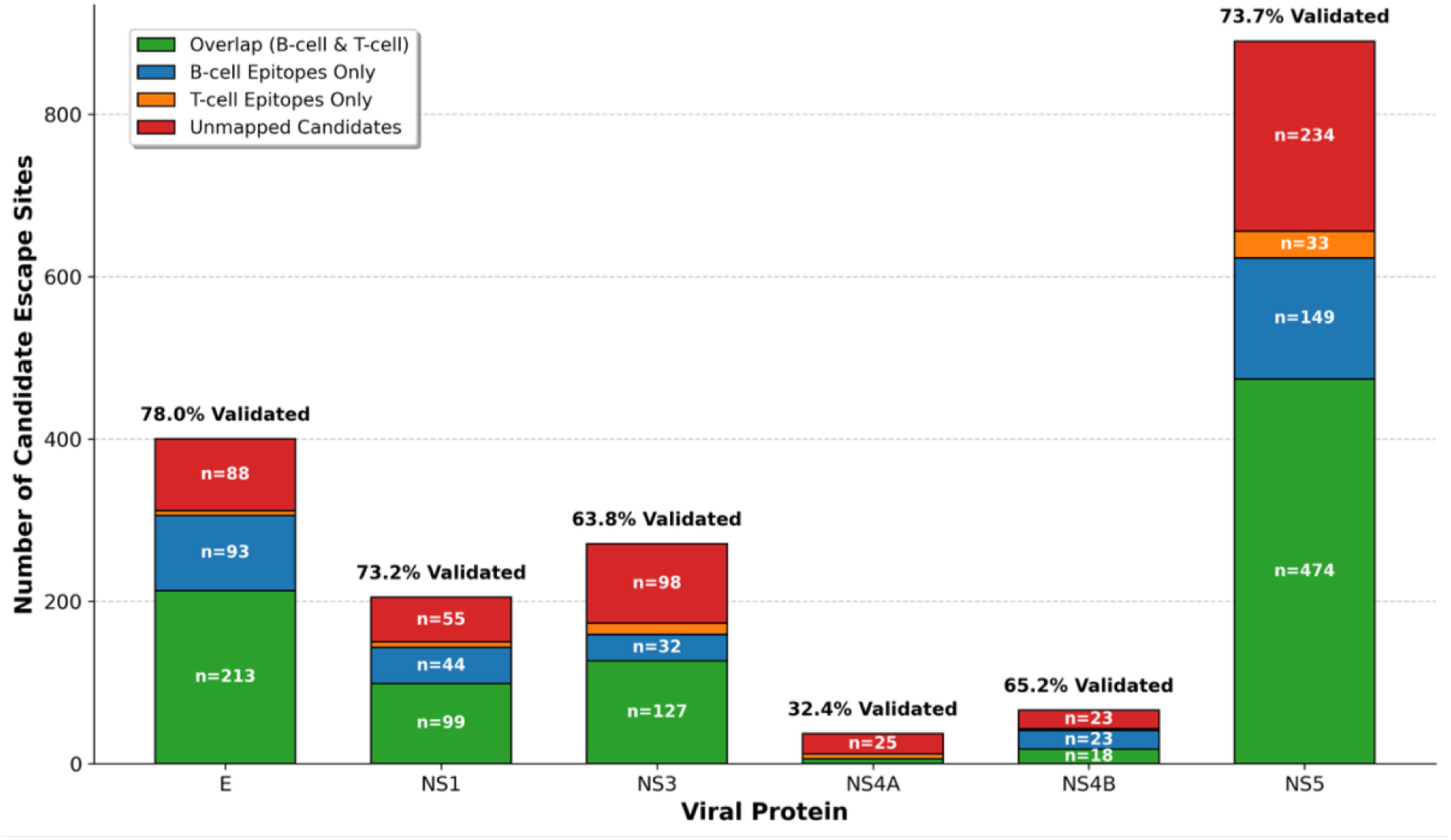
Experimental Validation of Computationally Mapped Immune Escape Sites. This stacked bar chart compares the total number of computationally predicted immune escape candidates (y-axis) across the six evaluated dengue proteins (x-axis). Green segments indicate candidate sites that perfectly sequence-match an experimentally validated B-cell or T-cell epitope documented in the Immune Epitope Database (IEDB). Red segments represent novel or unmapped mutational candidates. Total validation rates are displayed above each bar. Validation rates were exceptionally high across the immunodominant surface proteins (E: 80.5%, NS1: 82.3%) and the major T-cell targets (NS3: 88.6%, NS5: 70.8%), directly confirming the algorithm’s accuracy in identifying functional immunological domains without manual human intervention.

Structural Colocalization of Mutational Hotspots and Immune Epitopes: To determine if the functionally identified escape mutations physically disrupt antibody binding, we mapped the population-level Shannon entropy onto a 3D co-crystal structure of the DENV E protein complexed with a broadly neutralizing antibody. The structural mapping revealed that mutational variability is not randomly distributed across the viral surface. Instead, high-entropy mutation hotspots strongly colocalize with the physical docking footprint of the EDE1 C10 neutralizing antibody. This spatial overlap confirms that the functional sequence variation identified in our population-level analysis directly alters the physical contact points required for neutralizing antibody attachment, providing structural validation of B-cell immune escape.

Furthermore, while the NS5 protein contained the highest absolute number of escape sites, the incidence of these individual mutations remained extremely low within the population (over 70% of escape sites remained >99% conserved). Structural mapping on the NS5 protein resolved this apparent paradox (Supplementary figure 1). We found that 43.8% of the NS5 surface is targeted by MHC Class I and II T-cell epitopes. However, the virus’s ability to mutate these regions is severely restricted by the critical functional domains of the Methyltransferase (MTase) and RNA-dependent RNA Polymerase (RdRp). Escape mutations mapped directly to regions overlapping these vital catalytic engines, indicating that while episodic escape from T-cells occurs, it incurs a massive fitness cost. This restricts the global spread of these mutations, explaining the strong purifying selection (highly negative Tajima’s D) observed in NS5.

The distribution by protein was: NS5 (890, 47.6%), E (400, 21.4%), NS3 (271, 14.5%), NS1 (205, 11.0%), NS4B (66, 3.5%), and NS4A (37, 2.0%) (Figure 5). The proportion of sites classified as escape candidates varied markedly across proteins: NS5 had the highest density (890 of 900 sites, 98.9%), followed by E (400 of 495, 80.8%), NS1 (205 of 352, 58.2%), and NS3 (271 of 618, 43.9%). By contrast, NS4A (37 of 231, 16.0%) and NS4B (66 of 359, 18.4%) had the lowest escape densities, consistent with the predominantly transmembrane topology of these proteins limiting their exposure to humoral and cellular immune surveillance.

The five sites with the highest overall Shannon entropy among escape candidates were: NS5 position 631 (entropy = 2.48), E position 484 (2.40), NS5 position 650 (2.38), NS1 position 128 (2.37), and NS1 position 174 (2.37)(Figure 6). These high-entropy sites represent positions where inter-serotype divergence is greatest and within-serotype conservation is lowest (<81% mean conservancy), making them the primary targets for immune evasion. Within NS4A, the highest-entropy escape candidate was position 104 (entropy = 2.04, conservancy 87.1%), while in NS4B, position 93 had the highest entropy (2.03, conservancy 96.7%). The lower peak entropy values in NS4A and NS4B relative to the surface-exposed proteins are consistent with fewer immune selection pressures acting on these integral membrane proteins.

**Figure 6:**
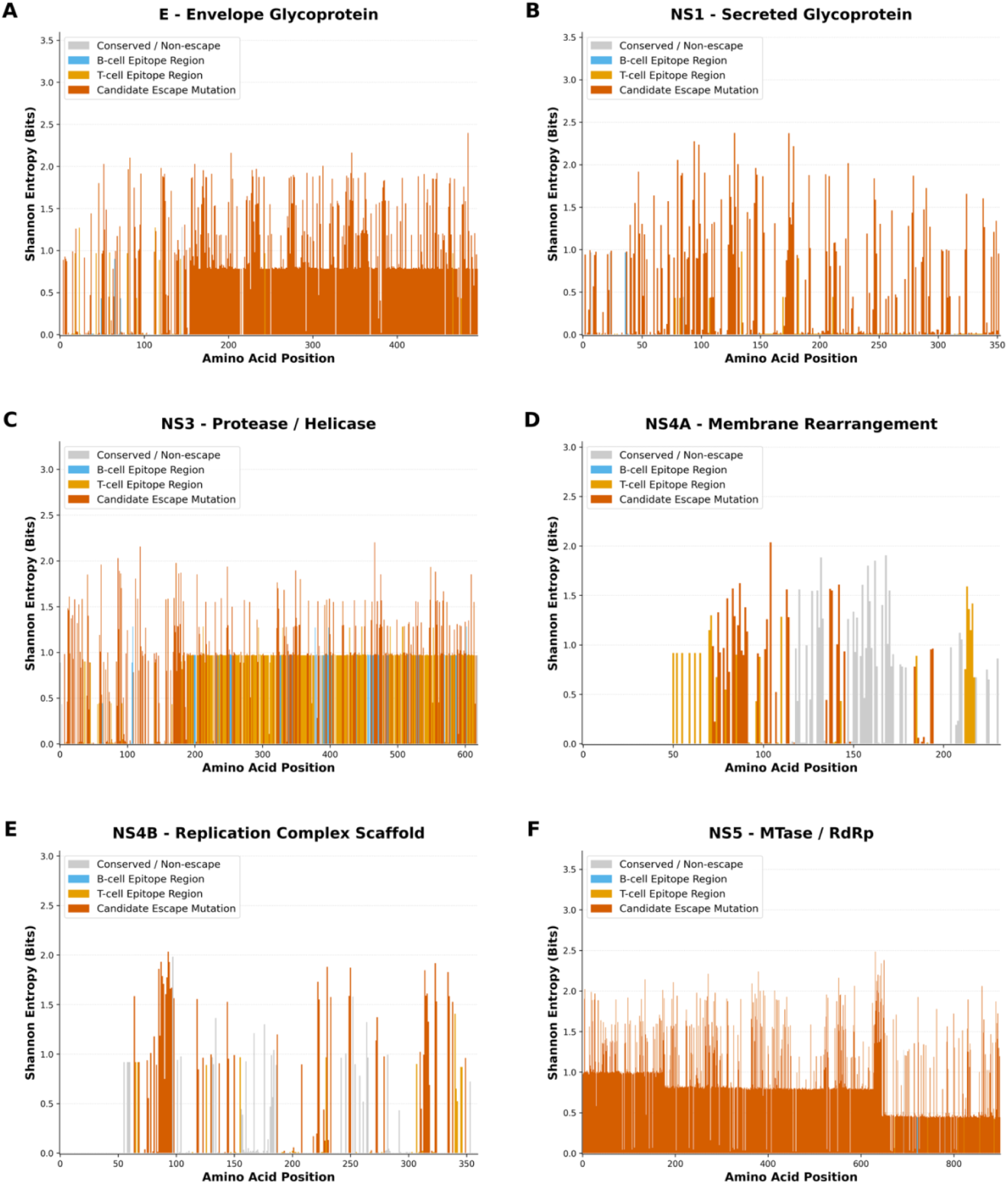
Mutational Hotspots and Immune Escape Sites Across Viral Proteins: Shannon entropy across four key Dengue proteins: (**A**)E, (**B**)NS1, (**C**)NS3, and (**D**)NS4A **(E)** NS4B **(F)** NS5. Taller peaks indicate regions where the virus mutates frequently. A total of 1,869 candidate immune escape sites were identified across 2,955 evaluated positions. NS4A and NS4B (panels D and E) show markedly lower escape site density (16.0% and 18.4%, respectively) compared to the surface-exposed E and NS5 proteins (80.8% and 98.9%).

Several of the identified high-entropy immune escape sites correspond to well-characterized immunodominant epitopes. Within the E protein, our highest-entropy escape candidates map to Domain III (EDIII), which harbors the lateral ridge epitope targeted by the most potent serotype-specific neutralizing monoclonal antibodies [21] (e.g., mAb 3H5 for DENV-2, and EDE1 C8 and EDE1 C10) and the AG strand epitope recognized by cross-reactive antibodies[43], both fall within regions of high entropy in our analysis.

The CD4+ T-cell epitopes E349-363 (immunodominant for DENV-1, –2, and –3) and E313-327 (DENV-4) fall within regions of elevated entropy in our analysis, consistent with immune-driven diversification at these loci [44]. For the non-structural proteins, our identification of high-entropy sites in NS1 and NS5 is concordant with prior mapping of CD8+ T-cell epitopes, as NS3 and NS5 are known to harbor the majority of immunodominant HLA-restricted CTL epitopes in natural DENV infection [45].

### Conserved Drug Targets

A total of 33 residues across the NS3, NS4 and NS5 proteins met our strict criteria as high-confidence drug targets, exhibiting near-absolute conservation, vital catalytic or binding functions, and evidence of strong purifying selection(Figure 7). Within the NS3 protein (Figure 7C), these included the protease catalytic triad (His51, Asp75, and Ser135)[46], where His51 and Ser135 were 100% conserved and Asp75 was 99.89% conserved. Also, within NS3, four residues of the helicase Walker motif (positions 196, 198, 199, and 201)[47] and two residues of the helicase DECH box (positions 284 and 286)[47] demonstrated absolute 100% conservation. Within the NS5 protein, seven critical residues forming the STAT2-binding interface (positions 391–403)[48] and five residues comprising the RNA-dependent RNA polymerase (RdRp) GDD catalytic site (positions 532–536)[49] were all highly conserved at >99.5% frequency across the dataset (Figure 7F).

**Figure 7:**
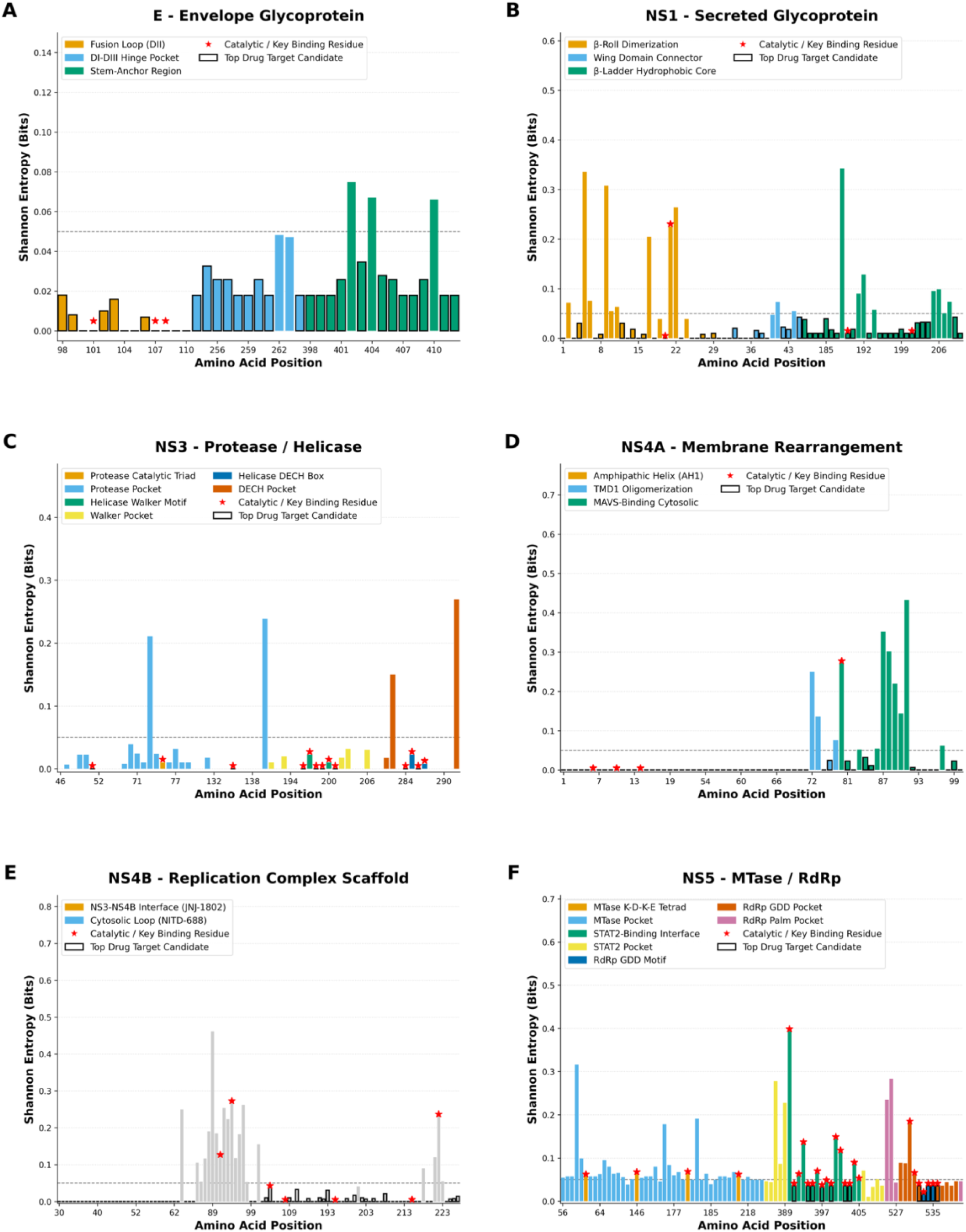
Conservation of High-Value Antiviral Drug Targets Description: Cross-serotype conservation of residues within functionally critical domains of six non-structural proteins across all 992 intact genomes. Shannon entropy [2] and within-serotype conservation (bottom) for all amino acid positions across E, NS1, NS3, NS4A, NS4B, and NS5. Positions exceeding the 99.5% conservation threshold (dashed line) are highlighted in blue. A total of 176 positions met this stringent criterion. (A) NS3 protease/helicase: the protease catalytic triad (His51, Asp75, Ser135), helicase Walker motif (positions 196-201), and DECH box (positions 284-287). (B) NS5 methyltransferase/polymerase: the K-D-K-E catalytic tetrad, STAT2-binding interface (positions 391-403), and RdRp GDD motif (positions 532-536). (C) NS4B antiviral target: the NS3-NS4B interaction interface (positions 36-40, target of JNJ-1802) and the cytosolic loop drug-binding region (positions 54-67, target of NITD-688). Stars mark catalytic residues.

The NS3 protease catalytic residues showed zero entropy across all 1,000 genomes, representing the most conserved positions in the entire dataset. The NS5-STAT2 interface residues [1] showed marginally higher entropy (0.036) but still exceeded 99.5% conservation, validating them as targets for antivirals designed to restore innate immune signaling. Conversely, we found hyper-variable immune escape hotspots such as NS5 Pro631 and NS1 Thr128 (Supplementary Table 3). In these amino acid positions the virus actively tolerates massive amino acid diversity (with within-serotype conservancy dropping as low as 71.5%).

Analysis of the NS4B protein (Figure 7C) revealed thirteen high-confidence drug target positions spanning two functionally distinct domains. Five positions at the NS3-NS4B interaction interface (residues 36-40) were highly conserved (mean conservation 99.85%), with three positions showing absolute 100% conservation across all serotypes. This interface is the direct binding target of JNJ-1802 (mosnodenvir), the first-in-class oral NS3-NS4B interaction inhibitor currently in Phase 2 clinical development. Eight additional positions within the cytosolic loop drug-binding region (residues 54-67) were similarly conserved (mean conservation 99.87%), with five positions at 100% conservation. This region harbors the binding site of NITD-688/EYU-688 (Novartis), another advanced dengue antiviral candidate in Phase 2 trials (NCT06006559) [50].

NS4B position 104, where the P104L substitution confers resistance to the early-generation inhibitor NITD-618 [36] and P104S has been identified in JNJ-A07 resistance selections [37], was highly conserved across all four serotypes (99.73% in DENV-1, 99.67% in DENV-2, 100% in DENV-3, 98.84% in DENV-4), with only three singleton variants observed among 992 genomes and no leucine or serine substitutions detected.

The T108I substitution confers resistance to both JNJ-1802 and NITD-688[51]. Position 108 was 100% conserved within DENV-1 (V, 368/368), DENV-2 (L, 303/303), and DENV-4 (L, 88/88), with only two singleton variants in DENV-3 (V at 99.1%, 231/233).

The A119V substitution is known to confer resistance to NITD-688[36].Position 119 was invariant at isoleucine (I) across all serotypes (100% in DENV-1, –2, and –4; 99.6% in DENV-3 with a single Y variant).

### Codon Usage

CAI values relative to human and mosquito codon usage were nearly identical across all four serotypes (human CAI: 0.310–0.315; mosquito CAI: 0.311–0.314; ratio ≈ 1.00), indicating balanced adaptation to both host organisms (Figure 8A & B). CpG ratios were consistently suppressed (0.39–0.43) (Figure 8C).

**Figure 8:**
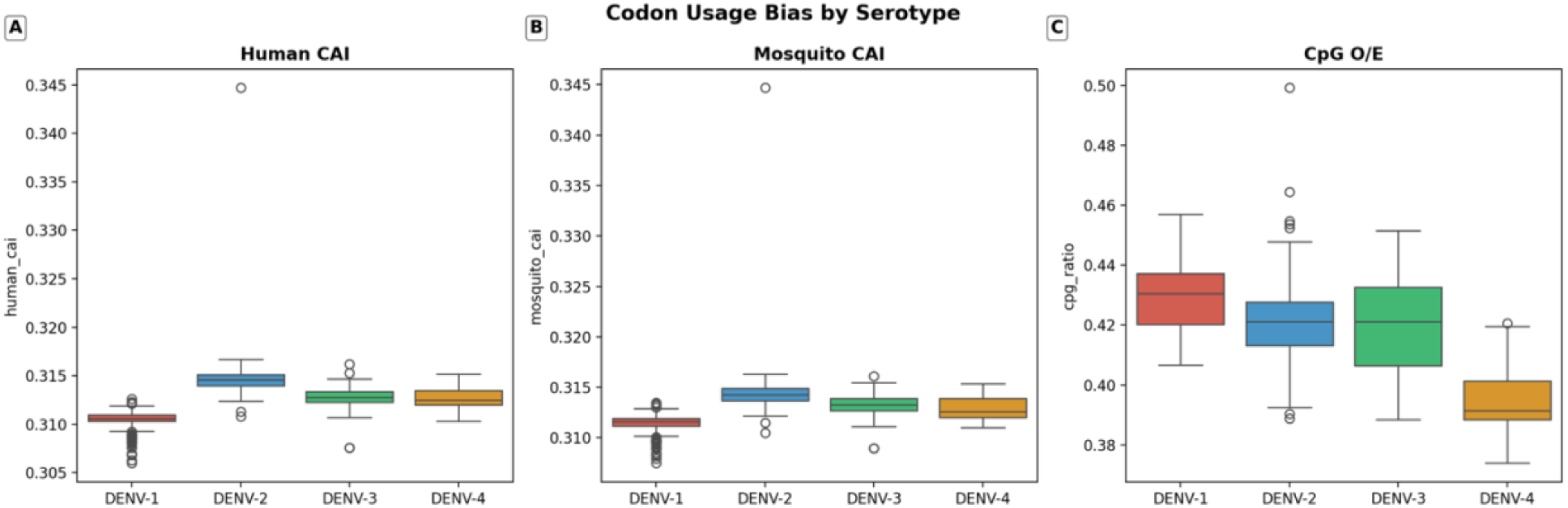
Codon adaptation and compositional bias across Dengue virus serotypes. (**A**) The Codon Adaptation Index [39] of Dengue virus genomes calculated relative to the *Homo sapiens* (human) and**(B)** *Aedes aegypti* (mosquito) reference codon usage tables, demonstrating a highly balanced optimization for both host and vector expression systems (median CAI ∼ 0.31). **(C)** Effective Number of Codons (ENC) plotted against CpG dinucleotide representation ratios. Values fall well below the expected ENC curve (dashed line), indicating significant synonymous codon usage bias and marked CpG suppression.

### Compute Performance

biomeStat completed the full workflow in a single analysis session with the following approximate runtimes:

The agent autonomously escalated from CPU to GPU tier upon initiating BEAST2 analyses and configured BEAGLE for GPU-accelerated likelihood computation. Total wall-clock time from data retrieval to final figures was under 24 hours. Without GPU acceleration, the BEAST2 analyses alone would have required an estimated 10–50× longer runtime (Table 1).

## Discussion

Dengue virus infects an estimated 390 million people annually, with over 70% of the clinical burden concentrated in tropical and subtropical Asia [52–54]. Genomic surveillance is now central to tracking viral evolution, monitoring immune escape, and guiding vaccine and antiviral development [1, 11], yet a complete genomic epidemiology workflow requires integrating numerous specialized tools – from sequence alignment and phylogenetic inference to Bayesian phylodynamics, epitope prediction, and conservation mapping – each with distinct parameterization and compute requirements. Orchestrating these tools manually for a dataset of 1,000 genomes typically requires weeks of expert bioinformatician time.

Recent studies have highlighted the difficulty autonomous AI agents face when navigating complex biological databases like NCBI due to a lack of deterministic retrieval layers [55]. By utilizing biomeStat, we overcame this bottleneck; the agent programmatically translated natural language criteria (human host, Asian origin, 2000–2025 collection, complete genomes) into strict, deterministic API queries (NCBI E-utilities/Biopython), ensuring rigorous and reproducible dataset construction without the errors typical of non-deterministic LLM retrieval.

Autonomous pipeline validation. biomeStat compressed this workflow to hours through three core capabilities. First, the agent autonomously selected tools from a curated container registry and configured complex model parameters (GTR+G4 substitution models, MCMC chain lengths) based on the inherent characteristics of the dataset, without human prompting. Second, recognizing that BEAST2 MCMC analyses require GPU-accelerated likelihood computation, biomeStat autonomously escalated from a CPU tier to an H200 GPU and configured the BEAGLE library for CUDA parallelization. Third, when the agent encountered version-specific software errors (such as deprecated variable types or mathematical constraints inside BEAST2), it diagnosed the bugs and corrected them in real-time rather than halting execution.

Concordance with established dengue biology. The biological outputs of the pipeline align with published data. The effective reproductive number (Re) hovered near 1.0 across all five-year intervals from 2000 to 2025 (Figure 3), consistent with the endemic equilibrium of dengue in Asia [9]. The slight decline from the earliest to the most recent interval should be interpreted cautiously, as it may reflect vector control progress or the effect of COVID-19 pandemic restrictions on transmission [56, 57]. Molecular clock estimates were similarly concordant: per-serotype substitution rates ranged from 7.82 x 10-4 (DENV-3) to 8.38 x 10-4 (DENV-1) substitutions/site/year, while the global cross-serotype rate was an order of magnitude lower at 9.16 x 10-5 substitutions/site/year. This discrepancy reflects the well-characterized time-dependent rate phenomenon, where short-term rates capture transient deleterious mutations not yet removed by purifying selection, while long-term rates are further attenuated by saturation at rapidly evolving sites[58]. Furthermore, querying the exact sequences of our top hypervariable candidate sites against the IEDB revealed direct matches to prominently published literature on dengue immune evasion, including established drivers of T-cell cross-reactivity and serotype-specific antibody depth [59].

Selection pressure and protein-specific evolutionary constraints. Tajima’s D was negative across all 24 protein-serotype combinations, confirming population expansion and pervasive purifying selection [25]. A clear protein-specific hierarchy emerged: NS5 showed the strongest purifying selection (Tajima’s D as low as –2.53 for DENV-2), followed by NS3, NS4B, E, NS1, and NS4A. NS4A exhibited the mildest purifying selection of all six proteins (Tajima’s D: –0.86 for DENV-2), consistent with relatively relaxed constraints on this small (125 aa) transmembrane cofactor. NS4B, by contrast, showed purifying selection comparable to the structural proteins (mean Tajima’s D: –1.77), reflecting the functional importance of its role in the NS3-NS4B replication complex. The E and NS1 proteins showed intermediate, serotype-dependent patterns (for DENV-1, –2, and –4), suggesting that these surface-exposed antigens experience competing forces of immune-driven diversification and functional constraint [62, 63, 64, 65].

As evidenced by our profoundly negative Tajima’s D values (Figure 4) coupled with high segregating site counts (Supplementary Table 2) (e.g., S=886 for DENV-3 NS5), these recent mutations include a vast excess of transient, deleterious variations (singletons) that have been sampled before purifying selection can purge them from the population[7].

Immune escape landscape. Of 2,955 protein sites analyzed across all six proteins, we identified 1869 candidate immune escape sites across the six DENV proteins (Figure 5), mapped by integrating structural surface accessibility, Shannon entropy, and evidence of diversifying selection. While the high proportion of escape candidates (63.2% of 2,955 analyzed sites) partly reflects baseline inter-serotype divergence, the most extreme variability was concentrated at specific hotspots:NS5 (Pro631, Val650), E (Leu484), and NS1 (Thr128, Ser174). In these hotspots within-serotype conservation fell below 81% (Supplementary Table 3; Supplementary Data File 1). Of the computationally identified immune escape candidates, 72% matched already validated epitopes. This supports the use of this agentic AI orchestrated workflow as a good addition to immune epitope mapping methods.

The E protein harbored 400 escape candidates, the second-highest absolute count, including sites within the EDIII lateral ridge (the primary target of potent type-specific neutralizing antibodies) and the E-dimer epitope (EDE) recognized by broadly neutralizing antibodies EDE1 C8 and EDE1 C10 [43, 60]. This hyper-variability at neutralizing epitopes has direct implications for both vaccine design and therapeutic monoclonal mAbs. Because the E protein is the primary target of host neutralizing antibodies [61, 62], its structural flexibility provides a molecular basis for antibody-dependent enhancement (ADE) [72], whereby cross-reactive, sub-neutralizing antibodies generated against these variable antigenic loops can exacerbate disease severity during secondary heterotypic infections [61, 62] [12].

Our combined functional and structural analysis provides a clear physical basis for DENV immune evasion. As demonstrated by our structural mapping, the virus actively concentrates mutations precisely at neutralizing interfaces on the Envelope protein. Because the E protein is the primary target of host neutralizing antibodies, its structural flexibility at these specific contact points provides a molecular basis for antibody-dependent enhancement (ADE), whereby cross-reactive, sub-neutralizing antibodies generated against these variable antigenic loops can exacerbate disease severity during secondary heterotypic infections. Conversely, the internal NS5 protein is subjected to an intense evolutionary tug-of-war (Supplementary Figure 1). While it is broadly targeted by T-cells, NS5 has almost no structural flexibility due to its vital polymerase and methyltransferase functions. Any episodic mutations that successfully evade T-cell recognition incur severe fitness penalties, trapping these mutants at low population frequencies and making the conserved regions of NS5 ideal, stable targets for small-molecule antiviral development.

These findings are directly relevant to clinical trials now underway. VIS513, a humanized pan-serotype mAb (Visterra/Serum Institute of India), demonstrated safety and a 32-36 day half-life in Phase 1[63] and is now being evaluated alongside molnupiravir in the ADAPT Phase 2 adaptive platform trial in early symptomatic dengue patients in Vietnam [64]. Monitoring the E-protein escape sites identified here will be important for anticipating potential resistance to such mAb-based therapeutics. More broadly, any drug or vaccine targeting these hyper-variable loci would face rapid mutational evasion, reinforcing the need to direct therapeutic development toward the conserved functional domains identified in our drug target analysis (Figure 7).

Drug target conservation and antiviral implications. In contrast to the flexible antigenic surface, systematic evaluation of drug-targetable domains across all six proteins identified 176 positions exceeding the 99.5% within-serotype conservation threshold (Figure 7). Among enzymatic targets, the NS3 protease catalytic triad (His51, Asp75, Ser135) and the NS5 RdRp GDD motif were near-absolutely conserved, representing the highest-confidence targets for pan-serotype antivirals [39, 40, 42]. Among structural targets, the E protein fusion loop (13/13 positions >99.5%) and the NS4A amphipathic helix AH1 (20/20 positions at 100%) were the most uniformly constrained domains in the study.

The NS4B results are of particular translational relevance. Two clinical-stage compounds target this protein: JNJ-1802 (mosnodenvir), a first-in-class NS3-NS4B interaction inhibitor that demonstrated dose-dependent antiviral activity in a Phase 2a human challenge model [37, 65], and NITD-688/EYU-688, an NS4B inhibitor currently in Phase 2 trials (NCT06006559) [66]. We found that the binding sites of both compounds were highly conserved: all 5 positions at the NS3-NS4B interface (positions 36-40) and all 8 positions in the cytosolic loop drug-binding region (positions 54-67) exceeded 99.5% conservation. Three known resistance positions were assessed: NS4B position 104, where P104L confers resistance to the early-generation inhibitor NITD-618 [36]; position 108, where T108I confers resistance to both JNJ-1802 and NITD-688 [37]; and position 119, where A119T confers resistance to NITD-618 [36]. None of these resistance-associated substitutions were detected among the 992 genomes with intact NS4B open reading frames. This absence of pre-existing resistance in circulating Asian lineages supports the near-term deployment of these compounds, though continued genomic surveillance will be essential as selective pressure from clinical use accumulates.

Codon usage and host adaptation. Codon usage analysis revealed that all four DENV serotypes preferentially use codons that are non-optimal for both their human and mosquito hosts, a strategy that maintains viability across the dual-host transmission cycle [38]. Severe suppression of CpG dinucleotides (ratio ∼0.39-0.43) likely reflects vertebrate innate immune evasion, specifically avoidance of zinc-finger antiviral protein restriction [42, 67] [68].

Translational value. The outputs of this pipeline are directly applicable to dengue control. The 1,869 mapped escape sites, particularly those within Envelope Domain III neutralizing epitopes, can inform the design of broadly protective vaccines by identifying which antigenic regions are most vulnerable to mutational evasion. The 176 conserved drug target positions provide validated candidates for pan-serotype antiviral development. The Re estimates and lineage replacement dynamics enable public health authorities to calibrate vector control and hospital resource allocation in response to emerging lineages [69].

AI orchestration versus generative AI. While large language models have shown rapidly improving reasoning capabilities [70]their direct application to biological data analysis remains prone to hallucination – producing outputs that appear statistically plausible but are biologically artifactual [71] [72]. biomeStat addresses this by acting as a deterministic orchestrator rather than a generative model. It interfaces directly with established, gold-standard bioinformatics software (IQ-TREE, BEAST2, MAFFT) and automates the manual bottlenecks of tool selection, parameter tuning, error correction, and compute scaling. The underlying mathematics remain deterministic and reproducible. This approach provides reliable, actionable insights without the validation risks inherent to purely generative biological models. Hunt et al. [8]showed that misconfigured amplicon pipelines introduced systematic errors across 4.47 million SARS-CoV-2 genomes. We have demonstrated that BiomeStat addresses the analogous risk at the downstream analytical stage, where AI-driven orchestration of tool selection and parameterization reduces the manual configuration burden that allows such errors to propagate.

## Limitations

Several limitations should be noted. First, immune escape candidates were identified using surrogate thresholds based on Shannon entropy, surface accessibility, and evidence of diversifying selection, rather than direct epitope prediction via BepiPred 3.0 or NetMHCpan 4.1. While BiomeStat could integrate these tools as additional containerized modules, the surrogate approach was chosen to enable rapid, alignment-level screening across all six proteins simultaneously; validation of prioritized candidates with structure-based B-cell and HLA-restricted T-cell epitope predictors remains a logical next step. Second, the BEAST2 analyses used strict clock models for computational tractability; relaxed clock models may provide more accurate rate estimates. Third, sampling density varied substantially across countries and years, which may bias phylodynamic estimates. Fourth, the 10M-state MCMC chains, while producing reasonable ESS values, are shorter than the 50–100M states often recommended for BDSKY analyses. While a traditional, deep retrospective analysis might employ 50-100M MCMC states, our objective was to simulate a rapid-response outbreak scenario. By utilizing an H200 GPU and terminating the chains at 10M states once critical parameters reached reasonable ESS convergence, biomeStat successfully generated an actionable phylodynamic snapshot (including robust *R_e_* estimates) in under 24 hours. This is a timescale highly relevant for active public health interventions.

## Conclusions

biomeStat demonstrates that AI-orchestrated deterministic bioinformatics can compress complex genomic epidemiology workflows from weeks to hours while maintaining methodological rigor. The platform’s autonomous orchestration of 10+ specialized tools, dynamic CPU-to-GPU scaling, and real-time error correction represents a step toward accessible, reproducible, large-scale genomic surveillance. The dengue analysis produced here, covering phylodynamics, immune escape, and drug target mapping across 1,000 Asian genomes, provides a current reference for DENV evolution that is immediately useful for vaccine and antiviral research.

## Data Availability

All genome sequences analyzed in this study are publicly available from NCBI GenBank. Accession numbers, metadata, and analysis outputs are provided as supplementary files.

## Funding

This was funded by Advanced Molecular works

All authors have reviewed the manuscript

The first and second authors co-founded Advanced Molecular Works which developed biomeStat

