## Supplementary Figure 1, Supplementary Table 1, Supplementary Table 2 for "biomeStat: Using Agentic AI for Scalable Genomic Epidemiology Demonstrated Through End-to-End Analysis of 1,000 Asian Dengue Virus Genomes"

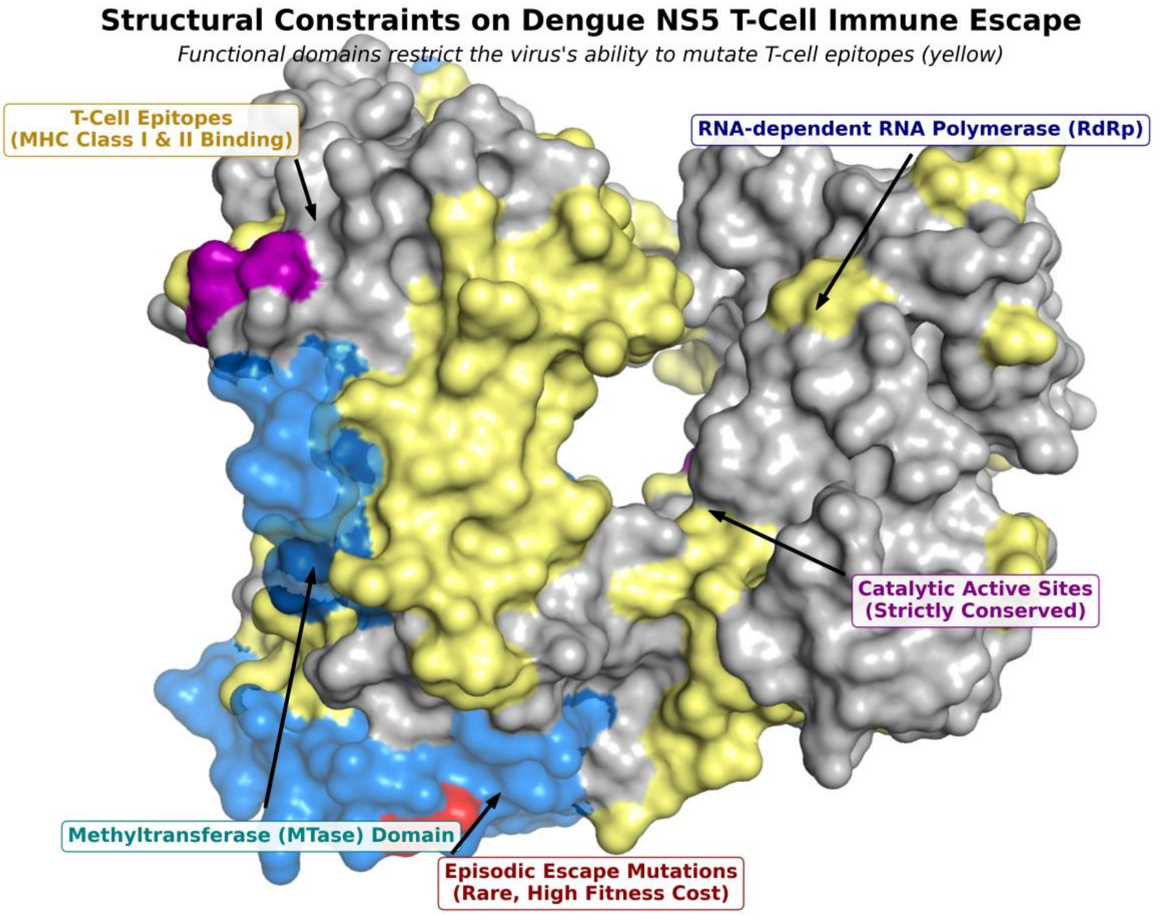


**Supplementary Figure 1. Structural constraints on episodic T-cell immune escape in the Dengue NS5 protein.** The 3D structure of the DENV NS5 protein (PDB ID: 6IZZ), comprising the Methyltransferase (MTase, highlighted in teal) and RNA-dependent RNA Polymerase (RdRp, highlighted in navy) domains. Regions of the viral surface predicted to bind host MHC Class I and II receptors (T-cell epitopes) are colored yellow, demonstrating that host cellular immunity targets a vast proportion (43.8%) of the NS5 surface. Strictly conserved catalytic active sites are indicated in purple. Episodic escape mutations (red), identified via population-level selection models (FUBAR/MEME), are highly restricted. The structural mapping demonstrates that mutational escape events occur directly within T-cell epitopes (yellow) but overlap with critical functional domains, imposing a severe fitness cost that restricts the global propagation of these variants and drives the strong purifying selection observed in NS5.

Supplementary Table2. Calculated Tajima's D values across the Envelope (E) and non-structural proteins (NS1, NS3, NS5) for all four Dengue serotypes. Strongly negative values indicate population expansion and/or purifying selection.

| Protein | DENV-1 | DENV-2 | DENV-3 | DENV-4 |
|---|---|---|---|---|
| E | -2.17 | -2.11 | -2.33 | -1.74 |
| NS1 | -1.85 | -1.77 | -2.47 | -1.21 |
| NS3 | -2.12 | -2.14 | -2.30 | -1.72 |
| NS4A | -1.677 | -0.856 | -1.667 | -1474 |
| NS4B | -2.052 | -1.504 | -1.761 | -1.749 |
| NS5 | -2.40 | -2.53 | -2.45 | -1.83 |

Supplementary Table 3. **Top Five Hyper-Variable Immune Escape Candidates Across the Dengue Virus Genome.** The five amino acid positions exhibiting the highest overall Shannon entropy among the 1,766 candidate immune escape sites identified in the dataset ($n=1,000$). Candidate sites were defined by their presence within a predicted B-cell or T-cell epitope, evidence of positive diversifying selection (via FUBAR/MEME), and elevated genetic variability. Overall, Shannon Entropy quantifies the global amino acid diversity at each position across all four serotypes. Mean Within-Serotype Conservancy reflects the average fraction of genomes retaining the consensus residue when calculated independently within each of the four serotype clades. The profoundly elevated entropy and depressed within-serotype conservancy (e.g., 71.5% at NS1-128) indicate that the virus actively tolerates and maintains extreme genetic variability at these specific antigenic loci, likely to evade host immune recognition.

| Protein | Position | Consensus Amino Acid | Overall Shannon Entropy | Mean Within-Serotype Conservancy |
|---|---|---|---|---|
| **NS5** | **631** | **P** (Proline) | 2.4847 | 0.8080 (80.8%) |
| **E** | **484** | **L** (Leucine) | 2.3965 | 0.8773 (87.7%) |
| **NS5** | **650** | **V** (Valine) | 2.3796 | 0.8637 (86.3%) |
| **NS1** | **128** | **T** (Threonine) | 2.3723 | 0.7153 (71.5%) |
| **NS1** | **174** | **S** (Serine) | 2.3699 | 0.8347 (83.4%) |
